## Supplementary Figures 1-3 for "Transposon-directed insertion-site sequencing (TraDIS) analysis of *Enterococcus faecium* using nanopore sequencing and a WebAssembly analysis platform"

**Table of contents:**

1. Supplementary Figure 1: TraDIS overview (page 2-3).
2. Supplementary Figure 2: TraDIS experimental set up (page 4).
3. Supplementary Figure 3: Growth curves of AUS0233Δ*tetM* (page 5).

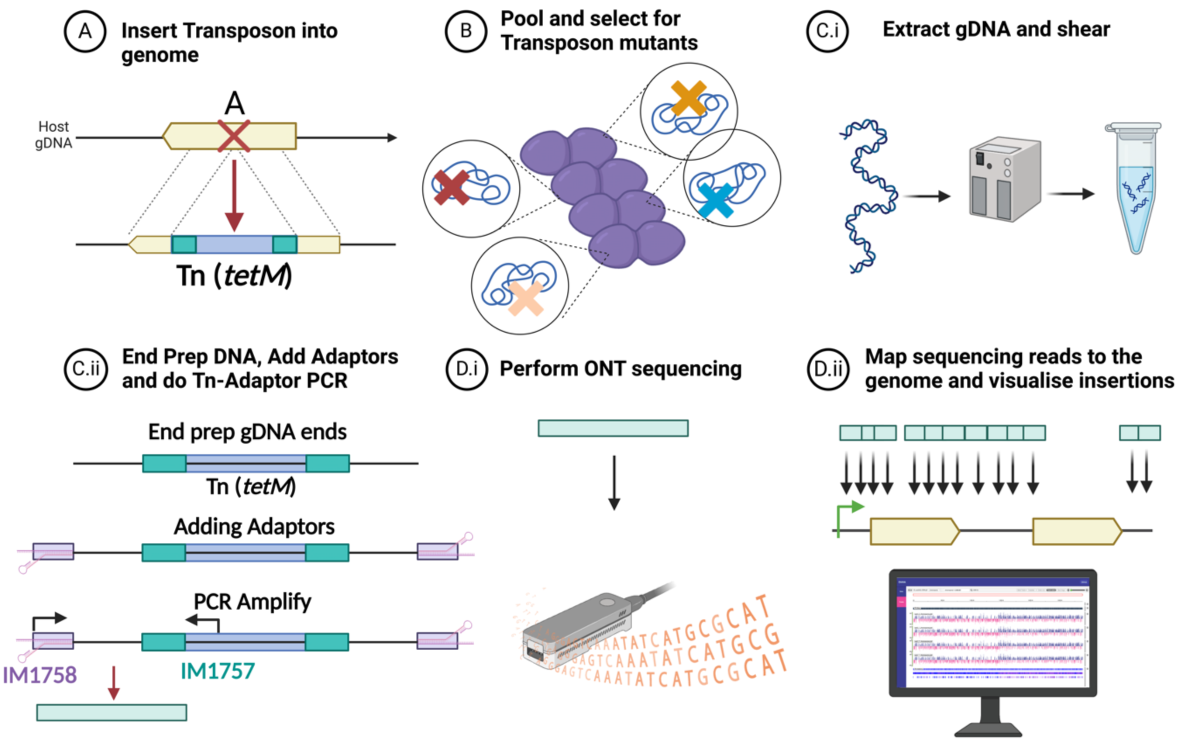


**Supplementary Figure 1. TraDIS overview**. *Construction and analysis of a TraDIS library. (A) Mutant library construction. The mariner transposon, flanked by inverted terminal repeats (green rectangles) is inserted into the host genome in a TA site within gene 'A' (yellow arrow). B) Library challenge. The coloured 'X's represent different transposon mutants, which are selected for on agar plates containing antibiotics to construct a pooled TraDIS library. The library can then be challenged, e.g. antibiotic exposure compared to an untreated control. (C) Transposon insertion site recovery. (C.i) TraDIS utilises a splinkerette-based PCR approach paired with high-throughput DNA sequencing to identify the flanking genomic region for each transposon insertion site. Genomic DNA is extracted, from both the challenge and control condition and then sheared by sonication to yield 0.5kb – 1.5kb DNA fragments. (C.ii) The splinkerette adaptor (purple rectangles) is ligated to end repaired genomic DNA. The transposon junction is captured by PCR amplification with the adaptor primer (IM1758) and transposon primer (IM1757). (D) Sequence analysis. (D.i) The PCR amplicon is barcoded*

*and then ONT sequenced using a R10.4.1 flow cell on a MinION. (D.ii) Processed reads from the ONT amplicons are mapped to a reference genome to identify insertion sites (black arrows). Subsequent bioinformatic analysis (Bio-Tradis) in Diana was conducted to identify conditionally essential genes, which contain significantly fewer insertions than non-essential genes, with data visualised as insertion plots.*

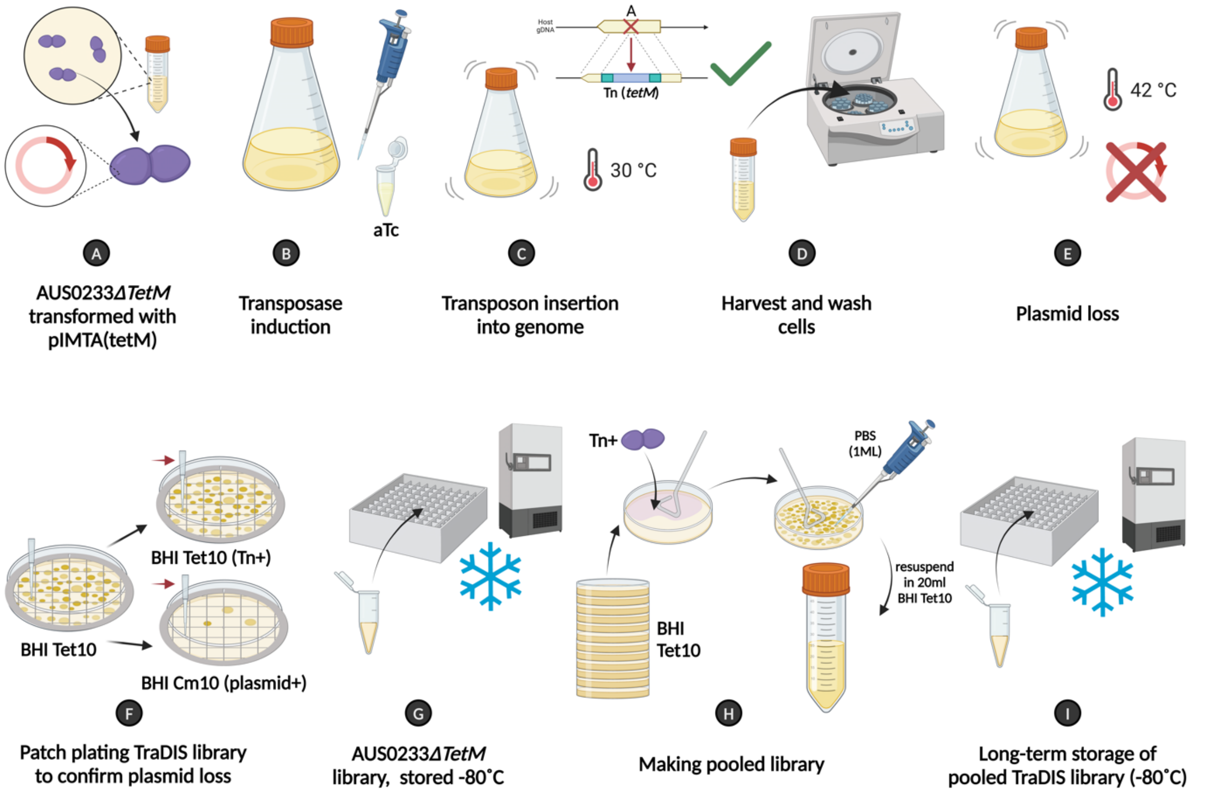


**Supplementary Figure 2. TraDIS experimental set up.** (A) *To generate a high density TraDIS library in E. faecium, strain AUS0233ΔtetM was transformed with pIMTA(tetM) at 30˚ C. (B) A colony was diluted in broth and the transposase was induced with 500ng/µl of aTc. (C) Transposition was carried out overnight. (D) The cells were washed to remove residual antibiotic. (E) Plasmid loss was stimulated by growth at 42˚ C. (F) Diluted aliquots of the TraDIS library were plated onto BHIA Tet 10µg/ml, with colonies then patch plated onto BHIA Tet 10µg/ml and BHIA Cm 10µg/ml to assess plasmid loss. (G) Individual aliquots of the TraDIS library were stored at -80˚ C. (H) Multiple separately induced TraDIS libraries were pooled to increase the number of unique transposon insertions. (I) Long term storage of pooled TraDIS library aliquots.*

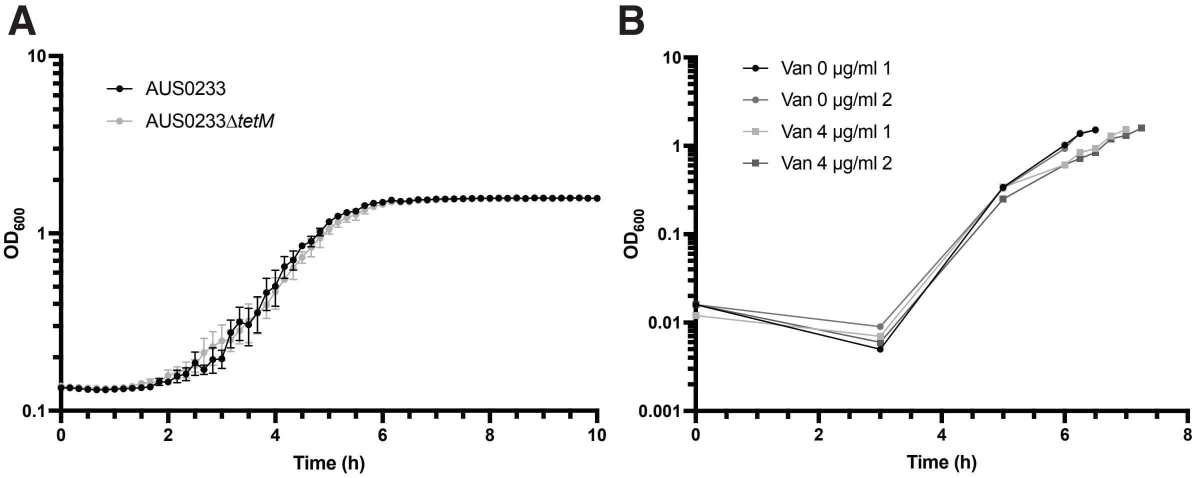


**Supplementary Figure 3. Growth curves of AUS0233Δ*tetM*.** *(A) Growth curve ($OD_{600nm}$) of AUS0233 and AUS0233ΔtetM strains in BHI. This data is representative of biological and technical triplicates, with the mean and standard deviation presented. (B) Growth curve of TraDIS library post-exposure to 0 or 4 µg/ml vancomycin at 0h. At $OD_{600}$ = 1.5, genomic DNA was extracted from the duplicate samples and processed for TraDIS.*
